# Small intestine lactobacilli growth promotion and immunomodulation in weaner pigs fed *Cyberlindnera jadinii* yeast high inclusion diet and exposed to enterotoxigenic *Escherichia coli* F4^+^: O149

**DOI:** 10.1101/2021.02.11.430732

**Authors:** S. Iakhno, S.S. Hellestveit, Ö.C.O. Umu, L.T. Bogevik, C.P. Åkesson, A.B. Göksu, C.McL. Press, L.T. Mydland, M. Øverland, H. Sørum

## Abstract

Enterotoxigenic *Escherichia coli* (ETEC) F4^+^: O149 is a causative agent for the development of post-weaning diarrhoea (PWD) in pigs that contributes to production losses. Yeast cell wall components used as a feed additive can modulate gut immunity and help protect animals from enteric infections. This work investigated how a novel yeast diet with high inclusion of yeast proteins (40% of crude protein) affected the course of ETEC mediated diarrhoea in weaner piglets from a farm with or without a history of post-weaning diarrhoea. We found that immune response to F4*ab* ETEC infection and appetite of the animals were altered by high inclusion *C. jadinii* yeast. The results indicate that the novel diet can support the diseased animals either directly through the effect of yeast beta-glucans and mannans or indirectly through the promotion of small intestine lactobacilli or both.

## 2 Introduction

Diarrhoea in neonatal and weaned piglets has been a concern to farmers due to the morbidity and mortality [1, 2]. The introduction of *E. coli* fimbrial vaccines [3] shifted the peak of diarrhoea from the neonatal and suckling period over to the weaning period where the mortality due to diarrhoea is lower [4]. An enterotoxigenic *Escherichia coli* (ETEC) of the O149 serotype has been incriminated in most of the post-weaning diarrhoea (PWD) cases contributing to production losses [2, 5–7]. This enteric pathogen acts via (I) the adhesion to small intestine enterocyte brush border with the help of receptor-specific fimbriae proteins F4 (K88) ( *ab*, *ac*, and *ad* variants) and (II) the production of toxins that induce enterocyte electrolyte/fluid imbalance hence watery diarrhoea. However, not all piglets are equally susceptible to ETEC. Some animals are immune to ETEC F4 *ab/ac* colonization due to an inherited trait that is thought to be linked to chromosome 13 of the pig [8]. A 74-kDa glycoprotein (GP74) was found to be key for ETEC adherence [9] but the genetic determinants encoding for this protein are not fully investigated [8, 10, 11]. Polymorphism in the *muc*4 gene was used as a basis for a DNA test to classify animals as either F4-adhesive or F4-non-adhesive [8]. Other candidate genes have been proposed as genetic determinants for the non-adhesive porcine phenotype [11]. The receptors for F4 *ab* fimbriae are found in the small intestine of newborn and weaned piglets [12] but not in older F4-adhesive animals [13]. While nursing piglets are protected from ETEC by maternal transfer of antibodies from vaccinated dams [3, 14], there are currently no measures available to protect piglets against ETEC-mediated diarrhoea after weaning (discussed in [15–17]). Modulation of the immune response against ETEC may be one such solution. Yeast cell wall components, mannans and beta-glucans proved potent immunomodulatory compounds. Fouhse and co-workers demonstrated that supplementation of yeast-derived mannans to weaner pigs positively affected jejunal villi architecture with corresponding changes in the gene expression profile [18]. The findings of Che et. al suggested that yeast mannans in feed could reduce systemic inflammation in pigs via suppression of lipopolysaccharide (LPS) induced TNF-alpha by alveolar macrophages [19]. Stuyven and colleagues reported protective effects of *Saccharomyces cerevisiae*, and *Sclerotium rolfsii* derived beta-glucans against ETEC F4^+^ with a reduction in pathogen shedding and F4-specific serum antibodies in weaner pigs [20]. Our previous work showed that feeding a strain of heat-inactivated *Cyberlindnera jadinii* yeast as a protein source changes the intestinal microbiota composition in weaner piglets [21]. Using cultivation and 16S *rRNA* gene metabarcoding sequencing techniques, we have shown that the yeast diet promoted the growth of small intestine lactobacilli. Beneficial immunomodulatory properties of intestinal lactobacilli are well documented ([22]; reviewed in [23]). These findings indicate that targeting the lactobacilli populations through diets can have an indirect impact on the host immune response.

Because beta-glucans and mannans are structural components of the yeast cell wall, and yeast replaced as much as 40% of the conventional proteins in the experimental diet, we hypothesized that *C. jadinii* yeast as a protein source can modulate the immune response towards ETEC F4^+^ and hence affect the course of PWD in weaner piglets.

To test the ability of a *C. jadinii* yeast diet to modify the course of PWD, we recruited piglets from two herds (with and without a history of PWD), primed them with either control or yeast-based a haemolytic F4*ab*^+^O149 *E. coli* isolated previously from the herd with the history of PWD. To diets where 40% of the protein was replaced with yeast, and orally challenged weaned piglets with a haemolytic F4*ab*^+^ O149 *E. coli* isolated previously from the herd with the history of PWD. To gain insights into the effects of yeast-derived feed, we compared gut microbial ecology metrics (diversity and composition), zootechnical performance, morphology and immunohistochemistry of gastrointestinal (GI) tract focusing on the ETEC F4^+^ intestinal colonization between the control and the yeast-fed piglet groups.

## 3 Results

### 3.1 General information

#### Post-weaning diarrhoea (PWD)

Of 68 piglets in the experiment, one animal from the control feeding group was euthanized *ad hoc* because of circulatory failure on d5 post-weaning (PW). There were no mortality cases due to the bacterial challenge throughout the experiment. Diarrhoea scores were higher for the first three days after the challenge in the piglets from the herd with no history of PWD (F4-naive herd) compared with those of the herd with the history of PWD (F4-immune herd) (Figure 2A).

**Figure 1:**
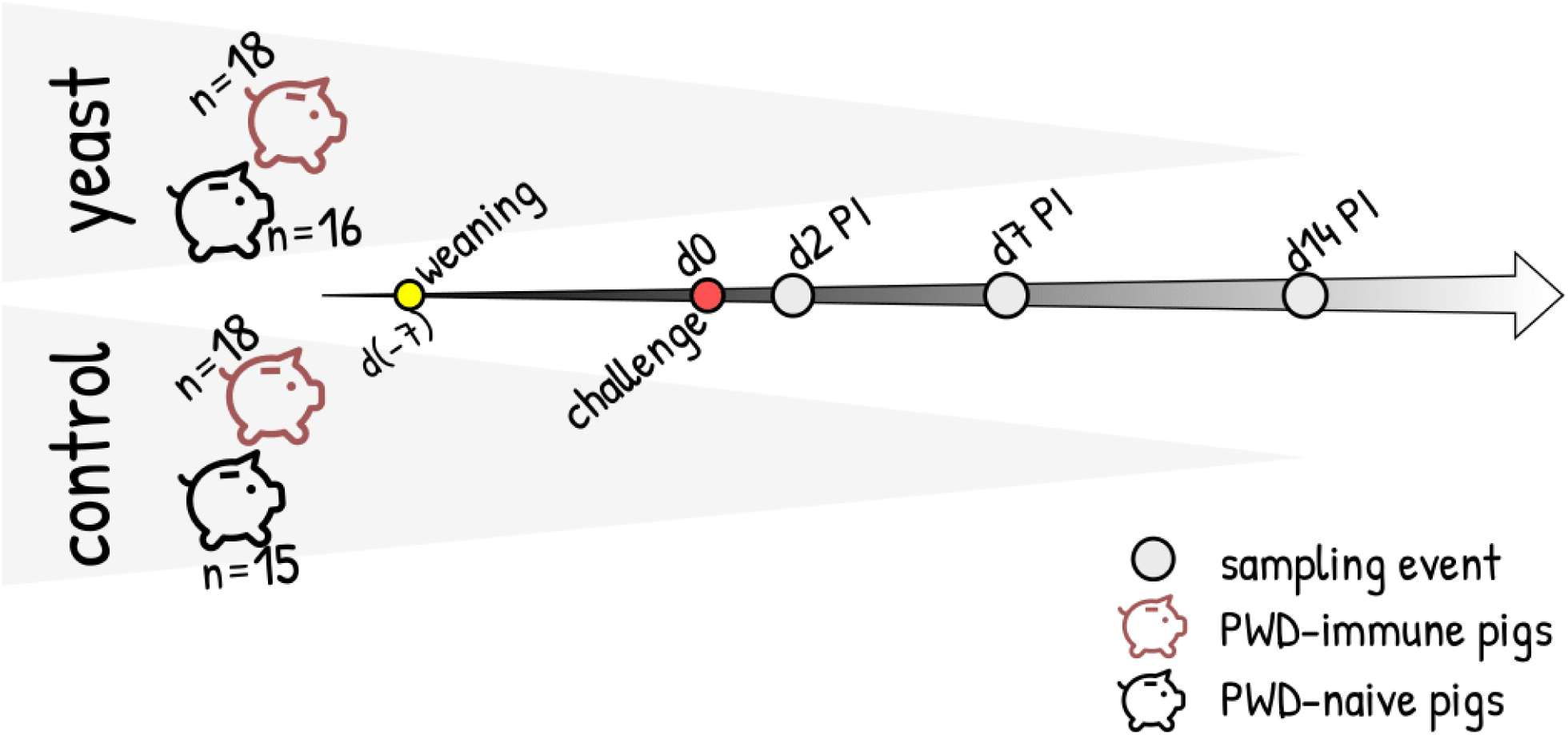
Overview of the experimental design

**Figure 2:**
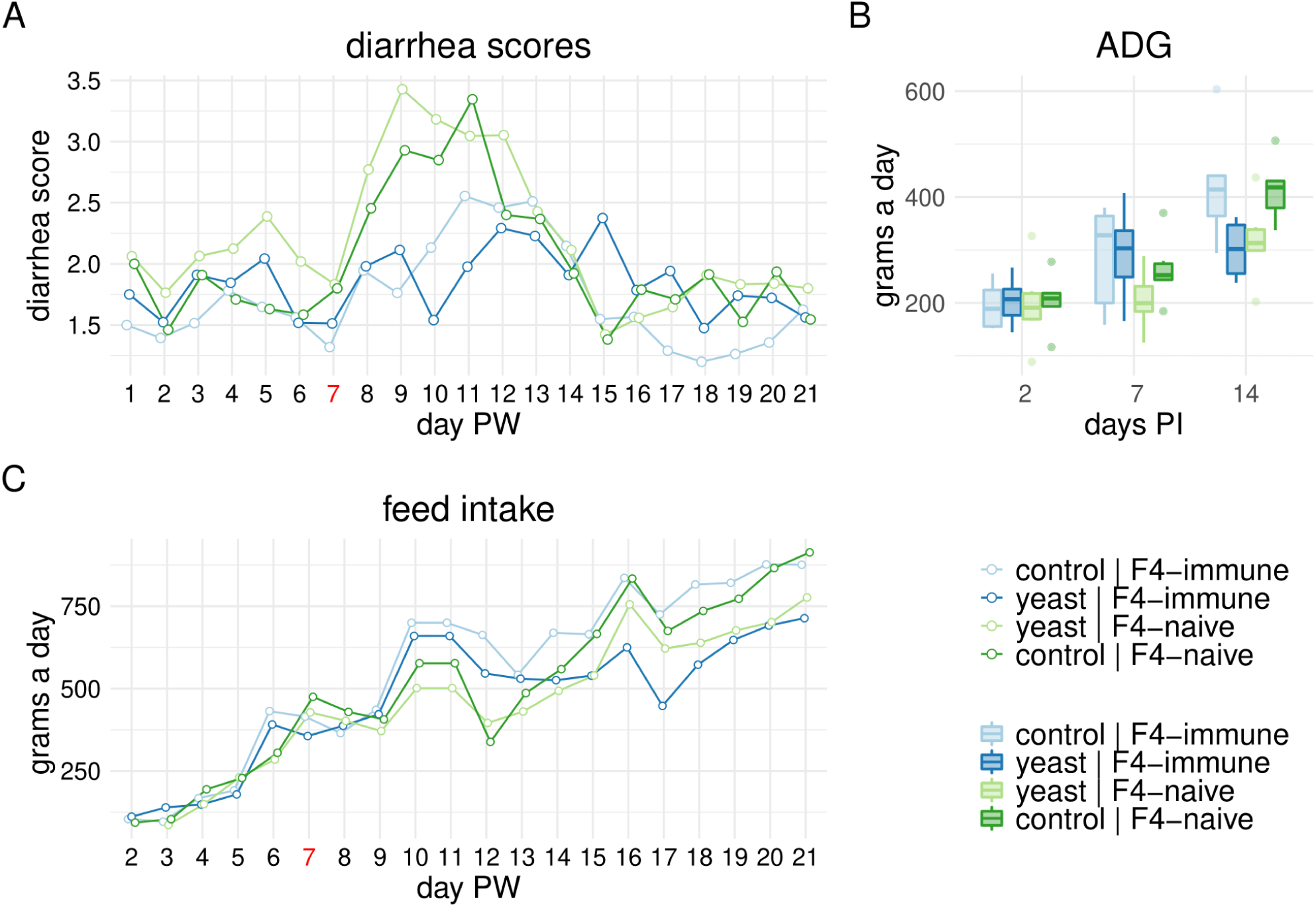
Diarrhoea scores and performance results. Panel A: Diarrhoea scores (pen level) across the experimental groups throughout the experiment. Day 7 post-weaning (coloured red) corresponds to the day the animals were orally challenged with ETEC F4^+^. Panel B: Distribution of the average daily gain (ADG) across the experimental groups at d2, d7, and d14 post-infection. Panel C: Daily feed intake across the experimental groups throughout the experiment. Day 7 post-weaning (coloured red) corresponds to the day the animals were orally challenged with ETEC F4^+^.

#### Average daily gain (ADG)

Average daily gain (ADG) was analysed by fitting the multiple regression model where “day”, “litter”, and “diet” were the predictor terms (d2 PI was excluded). The analysis revealed that the pigs fed the yeast-based diet tended to gain 62 g/day less weight than those fed the control diet (Figure 2B). The litter contribution to ADG estimate was as follows: litter3283, and litter3286 pigs tended to gain 125 g/day less than litter 3282 (p<0.00001); litter 3284 was gaining 86 g/day less than litter3282 (p=0.002); and litter3287 had 57 g/day greater ADG compared with that of the litter3283 (p=0.03).

#### Feed intake

The feed intake pattern (pen level) diverged between the herds from d3 PI to d5 PI with the F4-immune herd piglets eating more than those of the F4-naive herd. Within the herds, feed intake pattern showed that the control piglets ate more than the yeast fed piglets. From day 8 PI onwards, the effect of herd was less pronounced and changes in feed intake were attributed to the diet with the control group eating more feed than the yeast group (Figure 2C).

### 3.2 Immunohistochemistry

*F4 and CD3 in the ileum* **d2 PI** The proportion of the mucosa-associated ETEC F4^+^ per length of the ileum epithelium tended to be 5% greater in the pigs fed the yeast based diet than that of the pigs fed control diet (89% posterior probability)(Figure 3A). The piglets from the litter3288 had 10% less mucosa-associated ETEC F4^+^ per length of the ileum epithelium than that of the litter3282 (89% posterior probability) (not shown).

**Figure 3:**
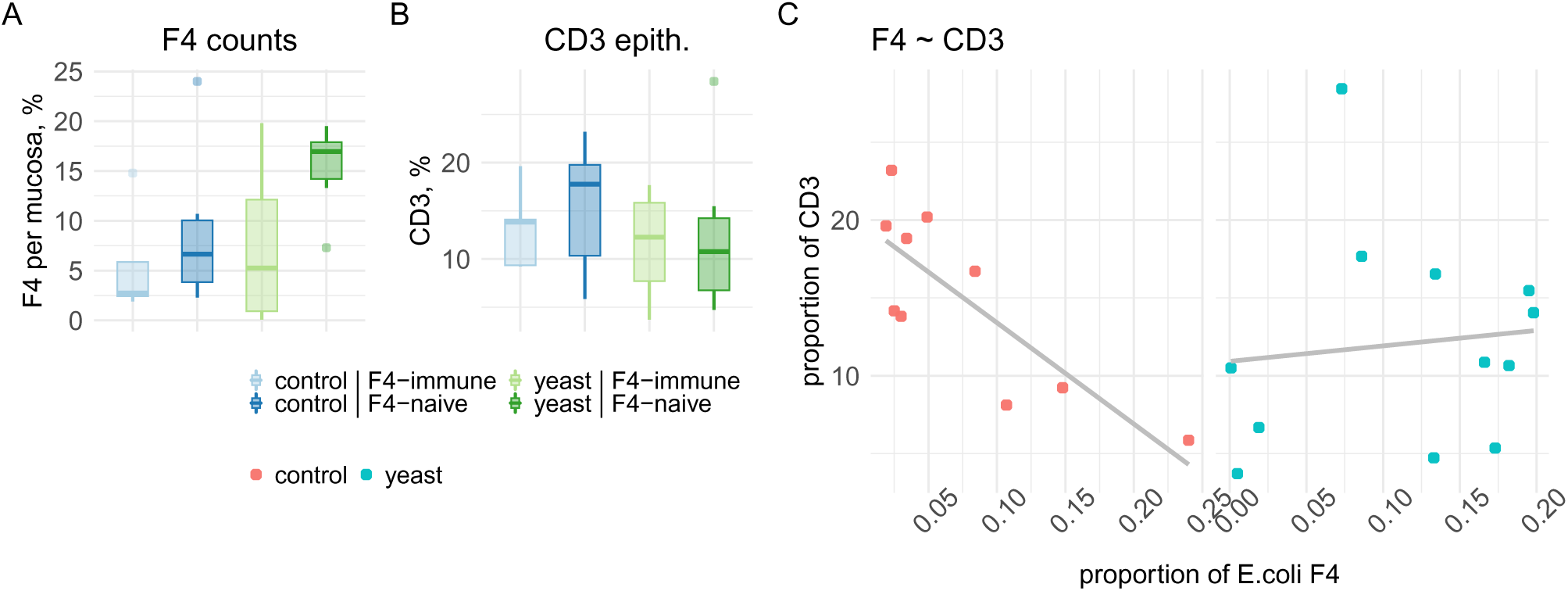
Immunohistochemistry results (d2 PI). Panel A: Distribution of the proportion of the mucosa-associated *E*. *coli* F4+ per mucosa section (lumen conten excluded) across the experimental groups on d2 PI. Panel B: Distribution of the proportion of IEL CD3+ cells in the epithelium across the experimental groups on d2 PI. Panel C: Correlation between the mucosa-associated F4+ *E*. *coli* and IEL CD3+ cells in the epithelium of control-fed (red dots) and yeast-fed (blue dots) piglets

At d7 PI, the prevalence of F4^+^ *E. coli* was lower in the ileum of the piglets fed both diets than that of d2 PI. Only two piglets in the yeast group had identifiable counts of F4^+^ adjacent to the epithelial surface compared with none of the control group. The remaining animals (n=16) were negative for the presence of F4^+^ *E. coli* in their ileum.

There was no clear relationship among the diet type, the litter, and the proportion of IEL CD3^+^ cells in the ileum epithelium of the pigs (Figure 3B). However, there was an inverse correlation between the proportion of mucosa-associated F4 antigen and the proportion of IEL CD3 populations in the ileum of the piglets fed the control diet at d2 PI (*rho*=-0.81, 95%CI upper = -0.25, 95%CI lower = -0.94) (Figure 3B). This relationship was not found in the yeast fed piglets (*rho*=0.1, 95%CI upper = 0.58, 95%CI lower = - 0.44) (Figure 3C).

### 3.3 Microbial ecology

#### 3.3.1 Sequencing results

Microbiota profiling was conducted on the ileum (n=63), caecum (n=67), and colon (n=66) digesta contents samples from pigs slaughtered on day 2, 7, and 14 PI (change to PW and same for the graph). Two sequencing runs produced a total of 58,045,034 raw reads. On average there were 71670 (SD=14239) reads per sample after filtering, denoising, and chimera removal (one sample with < 10,000 reads was deleted) (Supplementary Figure 10). Those reads were demultiplexed into 180, 856, and 906 unique amplicon sequence variants (ASVs) per ileum, caecum, and colon datasets, respectively (taxa not seen not more than once in 5% of a dataset were removed).

#### 2.3.2 Alpha diversity

Alpha microbial diversity comparison was made between the diet groups on day 2, 7, and 14 PI using the DivNet method to infer on the Shannon index. The ileum gut microbial communities of the yeast fed pigs were similar on the modelled Shannon index at d2 PI to those of the control diet. On d7 PI the ileum microbiomes of the yeast fed pigs showed a higher diversity than those of the control diet (Figure 4). This difference became more pronounced on d14 PI (Figure 4). As with the ileum, the microbial communities in the caecum of the yeast fed pigs were not different than those of the control at d2 PI. However, the caecal communities of the control diet-fed piglets were more diverse compared with those of the yeast diet (Figure 4).

**Figure 4:**
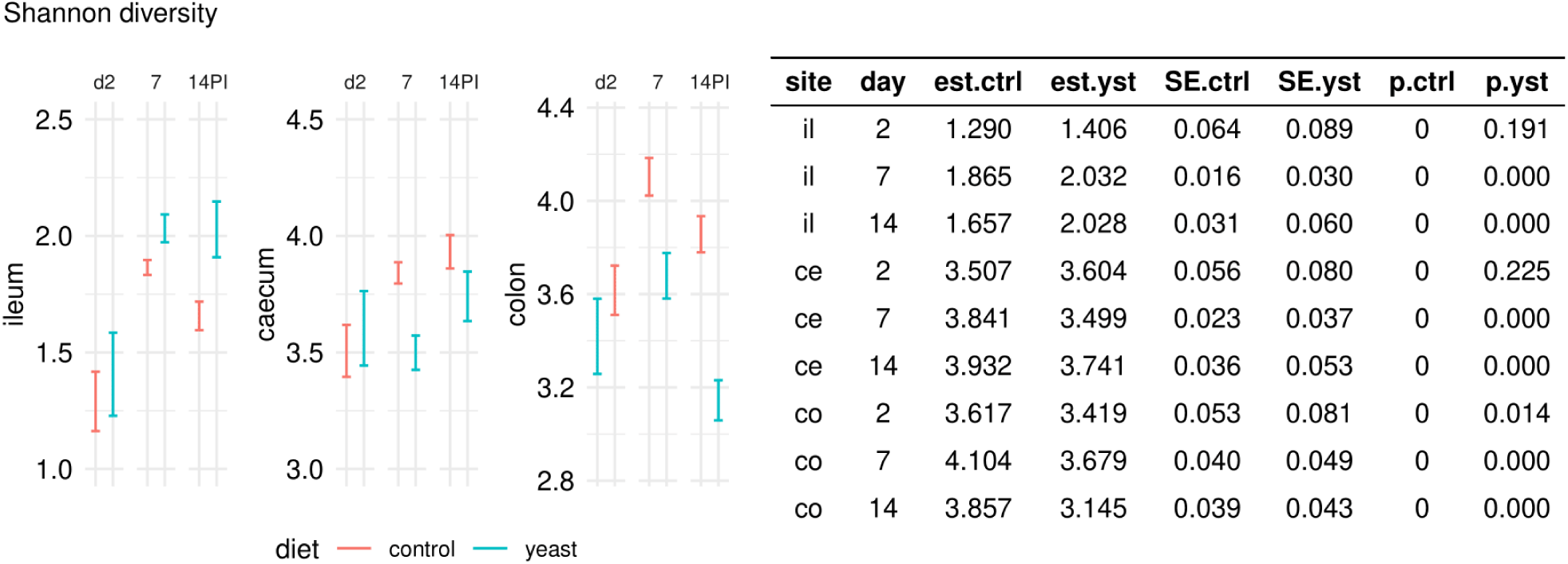
Alpha microbial diversity. Left: Estimates of DivNet inferred Shannon diversity indices with its uncertainty across gut sites (ileum, caecum, and colon), diets (control, yeast), and time (d2, d7, and d14 PI). The diet coloured intervals represent 4 standard errors (SE) (+2SE and -2SE around the estimate). Right: Summary of the DivNet statistical test for differences in the inferred Shannon diversity indices between the control and yeast diets: *site* shows the gut site microbiomes were derived from, *day* indicates the day post-infection when the samples were collected, *est*.*ctrl* and *est*.*yst* show the estimates of the Shannon index inferred by the model for the microbiomes of the pigs fed either the control or the yeast diets, respectively, *SE*.*ctrl* and *SE*.*yst* show the standard errors of the estimates of the Shannon index inferred by the model for the microbiomes of the pigs fed either the control or yeast diets, respectively, *p*.*ctrl* and *p*.*yst*, show the p-values derived from testing the difference in the Shannon diversity indices between the control and yeast groups, respectively

#### 3.3.3 Beta diversity

To study the impact of diets on beta microbial diversity in the intestines of ETEC challenged pigs, a multivariate model with permutations was fitted to the phylogeny-informed community data (see methods).

**day 2 PI** Although the **diet** was associated with the variance in the microbial communities on d2 PI across the **ileum, caecum, and colon** (R^2^ =9%), the **litter** (parental genetics) was a much **stronger predictor** of the variance in the respective microbiomes (R^2^ =38%) (Figure 5, Supplementary Figure 11).

**Figure 5:**
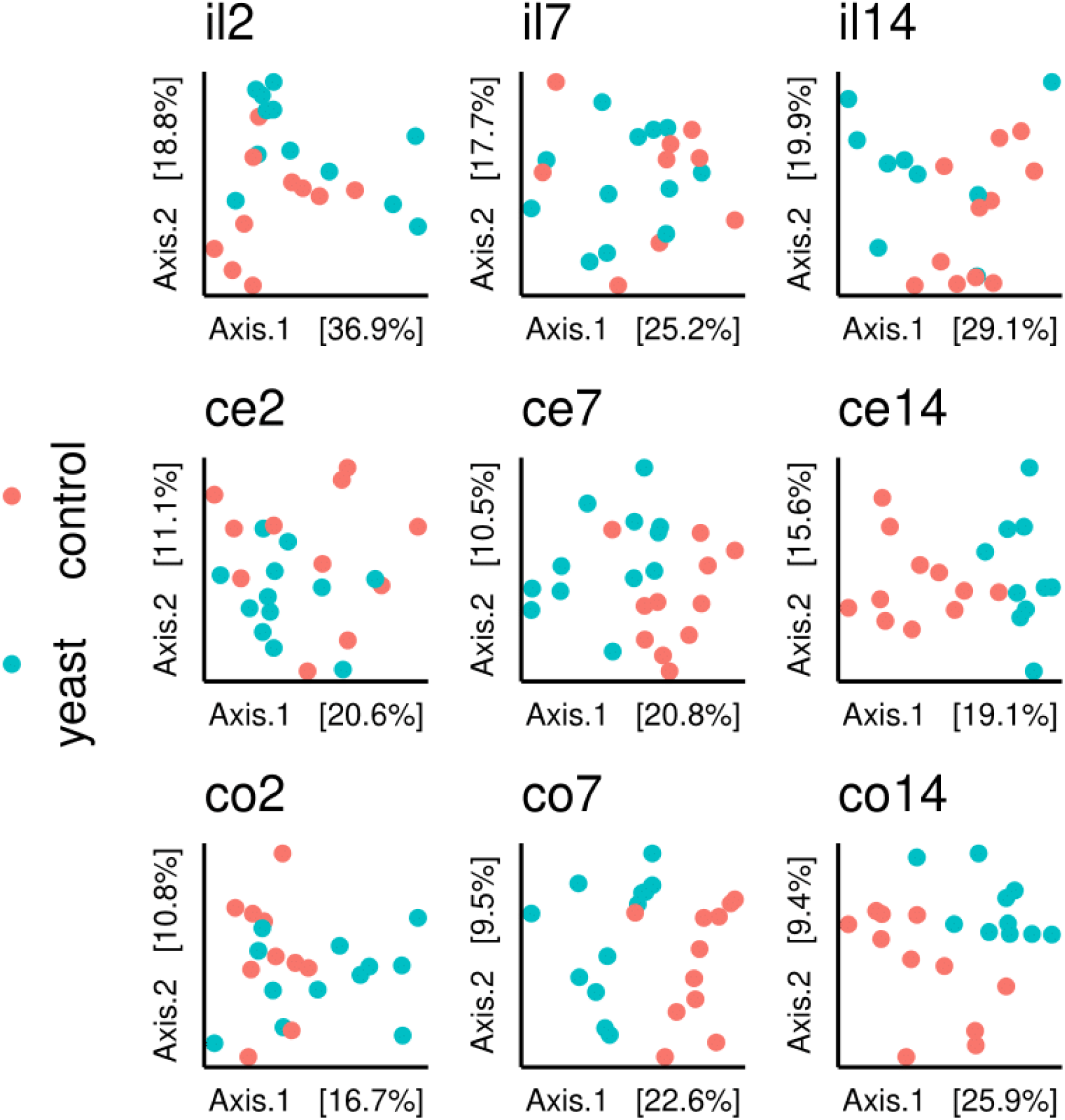
Beta microbial diversity. Principal coordinate analysis plot of the pig gut microbiotas coloured by diet (yeast, *blue*, control, *red*). The panel names designate distinct microbiomes across gut sites and time (ileum, *il*, caecum, *ce*, colon, *co* in combination with d2 PI, 2, d7 PI, 7, d14 PI, 14

**day 7 PI** The litter could predict 27.9% of the variance in the microbial data from the **ileum** of pigs sampled on d7 PI, while the diet was not a statistically significant predictor of the variance. The proportion of the variance in the microbial data explained by **diet increased** for the large intestine microbiomes at d7 PI (R =14.7%) compared with d2 PI. Reciprocally, the **litter** accounted for **less variance** of the unweighted Unifrac distances of the respective microbiomes (**caecum, colon** d7 PI) (R =24.2%) than that of d2 PI.

**day 14 PI** About the same amount of variance in the unweighted Unifrac distances was accounted by the **diet** across the **ileum, caecum, and colon** at d14 PI (R =14.2%), whereas the **litter was not** a statistically significant **predictor** of the variance at that time point.

#### 3.3.4 Differential abundance test

##### 3.3.4.1 Ileum

Two days after the challenge (d2 PI) there were more *Clostridia* class in the ileum microbiome of the control piglets compared with that of the yeast piglets. *Bacilli*, in contrast, were more predominant in the microbiome of the yeast fed piglets compared with that of the control (Figure 6). At a higher taxonomic resolution, a *Lactobacillus* cluster (sp. *reuteri*, *mucosae*, and *salivarius*) and *Streptococcus luteciae* were more predominant in the yeast microbiomes, while *Sarcina* and *Clostridium* sp. G060 were more predominant in the microbiomes of the control fed piglets.

**Figure 6:**
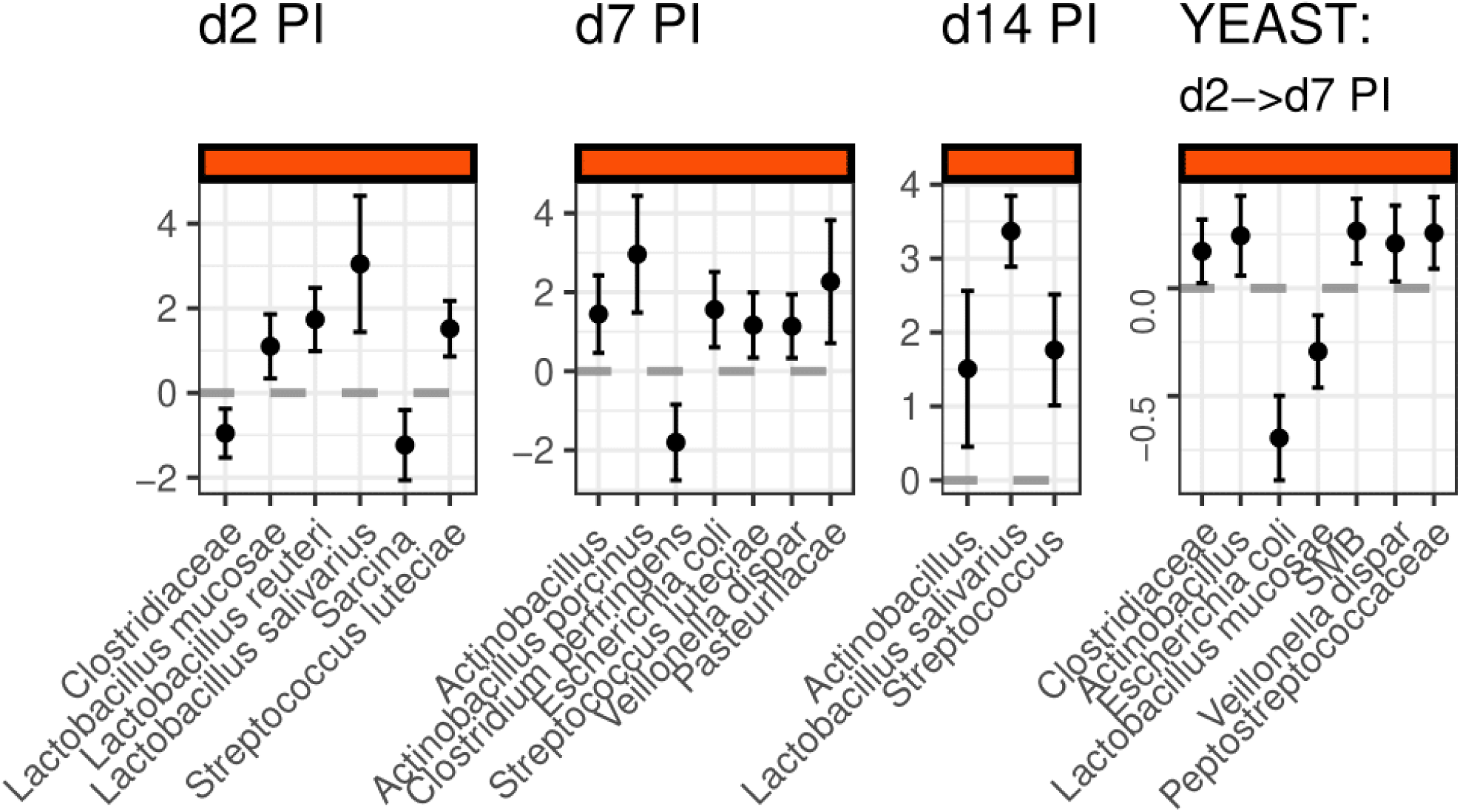
Differentially abundant taxa in the ileum (species level). The dots with the intervals represent the estimates of the beta-binomial regression model of the porcine faecal microbiomes along with its standard errors across d2-d14 PI; the positive estimates (above the grey dashed line, ”0”) indicate the taxa that are more predominant in microbiomes of the piglets fed the yeast diet compared with those fed the control diet. The *YEAST* panel shows differentially abundant taxa between the microbiomes of the yeast fed piglets at d2 and d7 PI; the positive estimates (above the grey dashed line, ”0”) indicate the taxa that are more predominant in microbiomes of the pigs on d7 PI in comparison with abundance on d2 PI

At d7 PI, the differential abundance of *Clostridia* and *Bacilli* bacterial classes was similar to the differential abundance at d2 PI (above). *Gammaproteobacteria* were more abundant in the microbiomes of the ileum of yeast-fed piglets compared to those of the control-fed piglets (Figure 6). More specifically, *E. coli*, *Streptococcus luteciae*, *Veilonella dispar*, *Actinobacillus* unclassified., *Actinobacillus porcinus*, and Pasteurellaceae ASVs were differentially abundant in the yeast-fed microbiomes of the ileum. Of note, *Clostridium perfringens* was more predominant in the ileum of the control diet-fed piglets.

At d14 PI, there again were more *Clostridia* class and less *Proteobacteria*, *Actinobacteria*, and *Gammaproteobacteria* bacterial classes in the control-fed ileum microbiomes compared with those of the yeast-fed piglets (Figure 6). At the family level, there were more Enterobacteriaceae, Streptococcaceae, Veillonellaceae, and Pasteurellaceae and less Clostridiaceae in the microbiomes of the yeast-fed piglets than those of the control piglets.

##### 3.3.4.1 Caecum

At d2 PI there were more *Streptococcus luteciae*, Paraprevotellaceae (CF231), and *Parabacteroides* taxa in the caecal microbiomes of yeast-fed piglets than in those of the control diet (Figure 7). At d7 PI the relative abundance of *Proteobacteria*, *Firmicutes*, *Deferribacteres*, *Actinobacteria*, and *Tenericutes* phyla were higher in the control fed piglet caecum microbiomes compared with those of the yeast (Figure 7). The only phylum that was more predominant in the yeast group caecum microbiomes than that of the control was *Bacteroidetes*. As many as 36 taxa were more predominant in the control fed piglet caecum microbiota compared with 2 taxa in that of the yeast (Figure 7).

**Figure 7:**
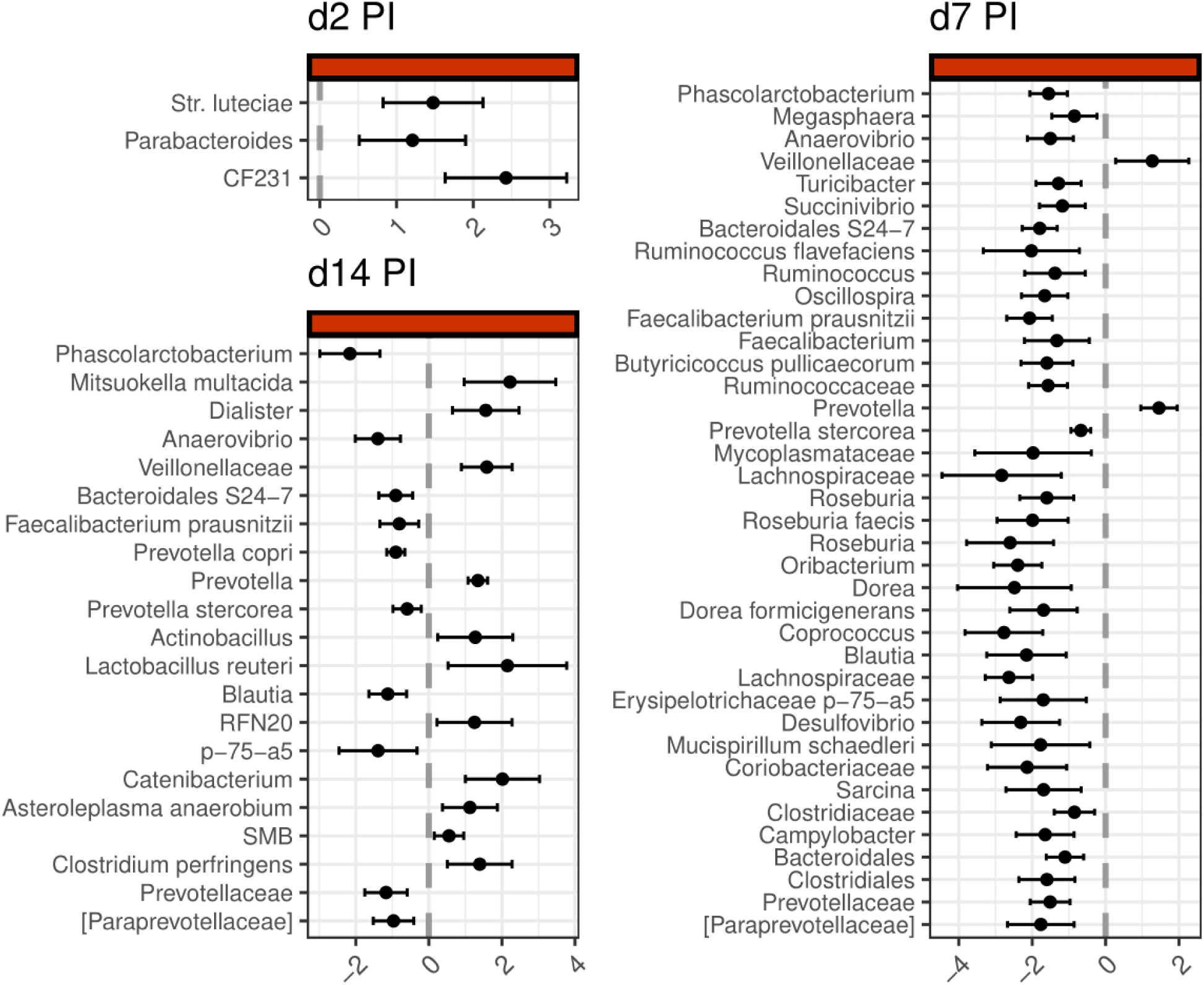
Differentially abundant taxa in the caecum (species level). The dots with the intervals represent the estimates of the beta-binomial regression model along with its standard errors across d2-d14 PI; the positive estimates (right of the grey dashed line) indicate the taxa that are more predominant in the microbiomes of yeast-fed piglets compared with those of the control-fed piglets

At d14 PI, the relative abundance of bacterial classes *Deltaproteobacteria* and *Erysipelotrichi* was differentially abundant in the yeast-fed piglet caecum microbiomes compared with those of the control-fed piglets. In contrast, *Epsilonproteobacteria* relative abundance was higher in the control-fed piglet caecum microbiomes compared with those of the yeast-fed piglets. At the species taxonomic level, there were 10 differentially abundant taxa in the control-fed caecum microbiomes compared with 11 of those in the yeast-fed piglets (Figure 7).

##### 3.3.4.1 Colon

At d2 PI, there more *Parabacteroides*, Paraprevotellaceae, Ruminococcaceae, and *Novispirillum* affiliated ASVs in the yeast fed piglet colon microbiomes than in those of the control-fed piglets. The relative abundances of *Campylobacter*, *Prevotella*, and *Desulfovibrio* were higher in the colon microbiomes of the control fed piglets compared with those of the yeast-fed piglets (Figure 8). At the species level of analysis, the relative abundances of *E. coli*, *L. johnsonii*, and *P. copri* were differentially abundant in the colon of control-fed piglets compared with those of the yeast-fed piglets (Figure 8).

**Figure 8:**
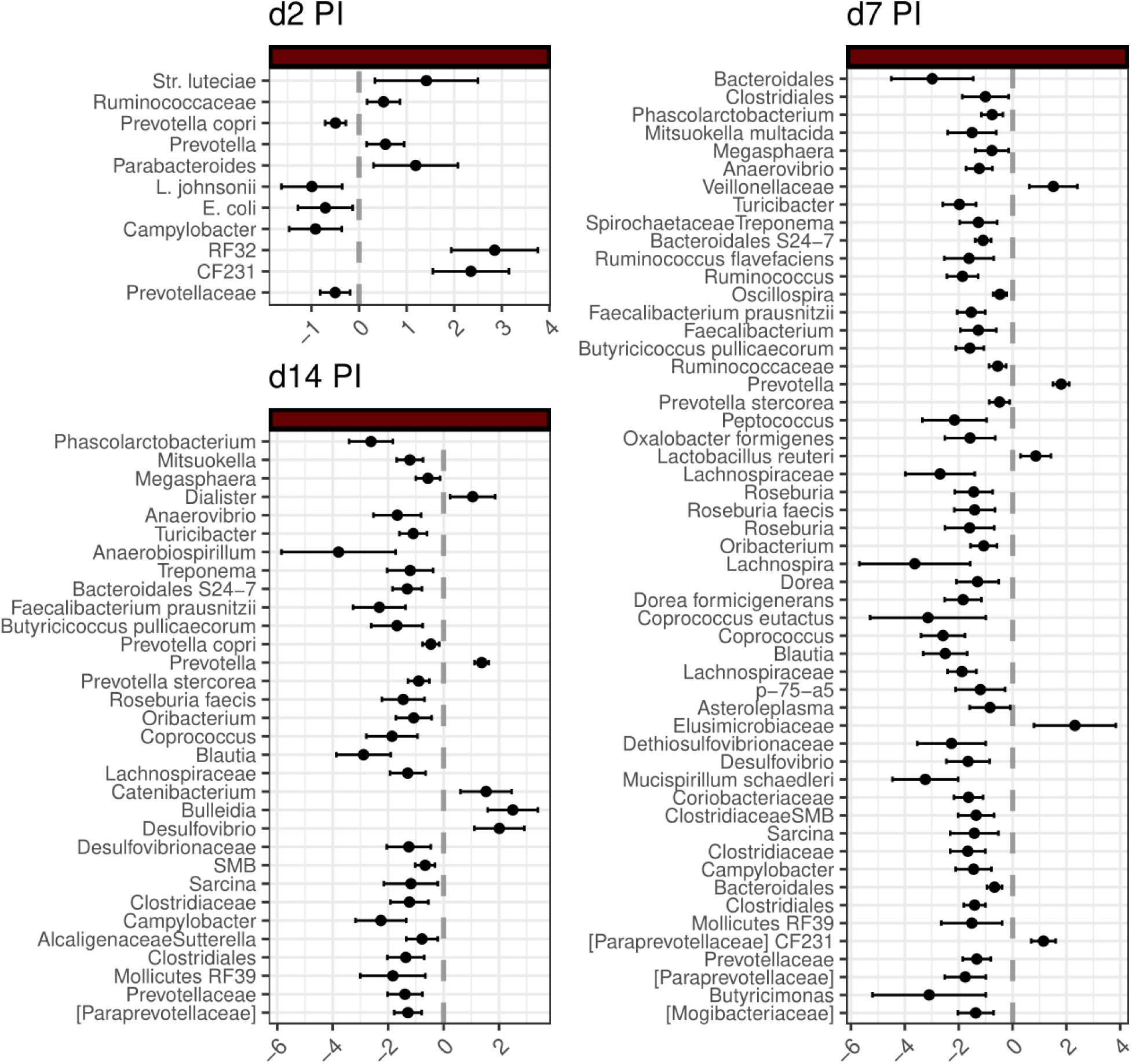
Differentially abundant taxa in the colon (species level)** The dots with the intervals represent the estimates of the beta-binomial regression model along with its standard errors across d2-d14 PI; the positive estimates (right of the grey dashed line) indicate the taxa that are more predominant in the microbiomes of yeast-fed piglets compared with those of the control-fed piglets

At d7 PI, the relative abundance of *Proteobacteria*, *Firmicutes*, *Spirochaetes*, *Deferribacteres*, *Actinobacteria*, and *Tenericutes* phyla was higher in the control fed piglet colon microbiomes compared with those of the yeast-fed piglets. *Bacteroidetes* and *Elusimicrobia* phyla were more predominant in the yeast-fed colon microbiomes than those of the control-fed piglets. At the species level, there were 48 differentially abundant ASVs in the colon microbiomes of the control-fed piglets and only 5 of those in the colon microbiomes of the yeast-fed piglets (Figure 8).

At d14 PI, the relative abundance of the bacterial phyla *Firmicutes* and *Tenericutes* was differentially abundant in the control-fed piglet colon microbiomes compared with those of the yeast-fed piglets. In contrast, *Bacteroidetes* phyla relative abundance was higher in the yeast-fed piglet colon microbiomes compared to those of the control-fed piglets. At the species level, there were 32 differentially abundant taxa in the control-fed piglet colon microbiomes compared with 5 of those in the yeast-fed piglet colon microbiomes (Figure 8).

### 3.4 Microbial network analysis

To characterize further the microbial communities that reside in the small intestine, microbial networks were recovered with the Sparse Inverse Covariance Estimation for Ecological Association Inference approach (SPIEC-EASI) algorithm (see material and methods).

The connectivity in the microbial communities of the ileum of the challenged pigs was sparse irrespective of time. Among the connected nodes, lactobacilli formed cliques more often than other phylotypes. Three members of the yeast fed pig microbiome lactobacilli, *L. mucosae*, *L. reuteri*, and *L. johnsonii*, were connected on d2 PI and d14 PI (Figure 9). *L. mucosae* which decreased in numbers in the digesta of the yeast-fed piglets (Figure 6), became disconnected from the lactobacilli clique on d7 PI (Figure 9). Lactobacilli of the control fed pig microbiomes formed bipartite cliques on d2 and d7 PI which consisted of *L. reuteri* and *L. johnsonii*. On d 14 PI theses two species were not connected (Figure 9)

**Figure 9:**
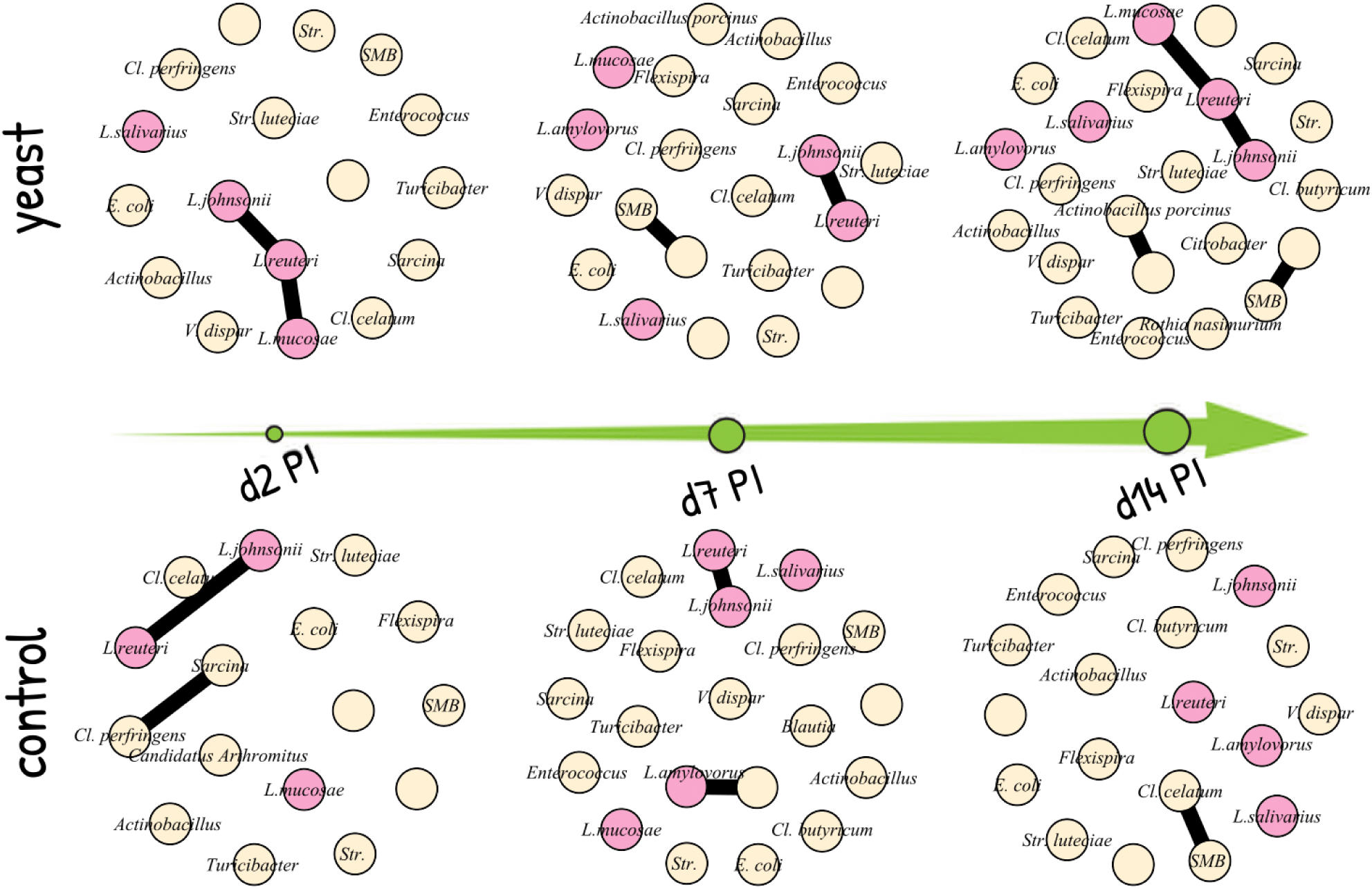
Microbial network of the ileum microbiomes across time and feeding groups. *Lactobacillus* genus is coloured pink, while other taxa are coloured in beige. The nodes (coloured circles) represent ASVs, while the black coloured lines represent connections between the nodes.

## 4 Discussion

This study investigated the impact of a novel yeast diet on weaner pig immunity assessed in the context of the intestinal microbiome and health parameters. The yeast diet contained beta-glucans and mannans as the structural components of yeast cell walls. Beta-glucans and mannans are believed to possess immunomodulatory properties when supplied to human and other mammals (reviewed in [24, 25]). In this study knowledge about the purity, quantity, and bioavailability of these compounds is limited. The heat deactivated *C. jadinii* yeast cells replaced 40% of crude proteins in the diet. The high dietary inclusion level suggests that large amounts of the immunomodulatory compounds were readily available to the weaned piglets through the experimental diet. A study by Håkenåsen et al. in healthy piglets fed a similar yeast diet demonstrated changes in the immune response in the small intestines by utilizing RNA sequencing analysis. Their findings featured an upregulation of immune signalling pathways, NF-kB and Toll-like receptors, already at d7 PW in the yeast-fed animals [26]. Lagos and co-workers showed that the *C. jadinii* yeast diet was associated with an increased CD3–/CD8+ cell population in distal jejunal lymph-nodes at d28 PW. However, the authors did not find this association in the blood [27].

In contrast to the studies of Håkenåsen et al. and Lagos et al., the present study employed an *E. coli* infection model to elicit changes in the immune response that are attributable to the yeast diet and were not evident in the healthy animal experiments. The choice of the challenge strain (F4ab) used in this study was guided by the relevance of that pathotype for the Norwegian and European swine industry [2, 6, 28]. Once established in a pig farm, the pathogen can remain in the environment for a long time and is hard to eradicate [1, 29]. Another important aspect of this bacterium is that suckling piglets are mostly immune to the infection through colostrum and milk from vaccinated mothers. Sow vaccination shifts the adhesive *E. coli* disease onset to the post-weaning period where piglet mortality due to PWD is lower compared with that of neonates [3, 4]. The reduced growth of the animals due to PWD, however, may be relevant for the industry. From the experimental point of view, this infection model was an appropriate replication of the field disease as the induced infection caused no mortality.

The degree of adhesiveness of F4^+^ *E.coli* to porcine enterocytes and subsequently the rate of bacterial colonisation is determined by the genetic constitution of the pigs. One such genetic determinants is an SNP located in the *muc4* gene of porcine chromosome 13. Others have suggested that additional SNP candidates are implicated in F4 susceptibility adhesion porcine phenotypes [11]. To our knowledge, the only DNA based assay that can discriminate between the adhesive and non-adhesive porcine phenotypes is the one developed by Jørgensen and colleagues [8]. The present study involved two principally distinct herds: one with a history of PWD (F4-immune) and another without a history of PWD (F4-naive). The F4-immune phenotype of the pig herds was supported by DNA testing. There were 19 animals in the F4-immune herd that had a mutant allele within the *muc4* gene compared with none in the F4-naive herd. Our observations of diarrhoea severity due to F4 *E. coli* supported the genotyping results related to F4 susceptibility. The diarrhoea scores were higher in the F4 naive herd piglets from d1 PI to d3 PI. This time-window corresponds to the classical development of PWD [20, 30]. The faecal scores in the F4-immune herd were only slightly elevated post-infection. Feed intake figures also highlighted a lower severity of PWD in the F4-immune herd than that in the F4-naive herd. After the acute phase of the ETEC infection, on d4 PI, the F4-immune piglets were eating more and gaining more weight compared with the F4-naive piglets. One of the key findings in the present study was that the yeast-fed piglets were eating less and subsequently gaining less weight d14 PI than the control-fed piglets. Unlike the figures at d7 PI, the effect of F4 susceptibility on the feed intake and ADG was not pronounced. These findings contrast with previous studies in healthy piglets where feed intake was comparable between yeast-fed and control-fed pigs [26, 31].

The implications of appetite loss in yeast-fed animals challenged with a pathogen are unclear. To our knowledge, PWD-affected piglets recover well, and there was no production loss due to the disease on the farm with a history of PWD (the National litter recording system, “Ingris”). It has been proposed that reduced appetite is an adaptation trait which, in concert with the immune response, helps mammals survive enteric infections [32]. Murray and colleagues demonstrated that food avoidance in mice infected with *Listeria monocytogenes* resulted in 50% less mortality compared with the infected force-fed mice [32]. Wang and co-workers [33] obtained similar results by reproducing the experiment by Murray and colleagues [32]. The listeriosis and colibacillosis infection models are not directly comparable concerning the mortality/morbidity rates. The design of this study precludes us from making assumptions on how herds without a history of PWD would fare after being exposed to PWD. However, here we can speculate that the development of appetite loss in the yeast-fed piglets might render pigs more robust against possible subsequent infectious stressor. A longitudinal study design, or a field trial, is essential to address this research question.

While changes in appetite were observed towards the end of the experiment, changes in the distribution of immune cell populations were already visible at d2 PI. There was an inverse relationship between the intraepithelial CD3 populations located in the ileum and the degree of F4^+^ *E. coli* colonisation in the control-fed piglets. In contrast, this relationship was not present in the yeast-fed piglets. This finding suggests that the yeast diet enabled intraepithelial T cell populations to persist in the presence of high levels of mucosa-associated F4^+^ *E. coli*.

Our results corroborate and elaborate on the findings of differences in the immune gene expression in the porcine small intestine reported by Håkenåsen et al. [26]. These investigators demonstrated that on day 7 after the introduction of yeast-based feed, several immune system pathways, including Toll-like receptor and NF-kappaB signalling pathways, were enriched in the small intestine of the animals. High inclusion levels of immunomodulatory yeast compounds in diets likely stimulates small intestine immunity.

It is our speculation that the immune system was (I) modulated prior to the infection either by the immunogenic compounds of the yeast cell walls or shifts in small intestine microbial communities or both and then (II) exposed to antigenic stimuli due to the ETEC infection. This speculation is encouraged by our observations of higher counts of F4^+^ *E. coli* in the F4-naive herd compared with those of F4-immune herd on the yeast diet. In other words, the growth of intestinal ETEC was suppressed in the pigs from the herd with a history of PWD.

These findings indicate the presence of an effect of the yeast diet on the local immune response and, later, on appetite. Hoytema van Konijnenburg et al. using a murine model showed that intestinal intraepithelial lymphocytes (IELs) movements within the epithelium are antigen-specific[34]. The authors demonstrated using live imaging that the IELs increased their motility within the epithelial cell layer (“flossing”) when exposed to *Salmonella enterica* antigens. Also, they found that in the absence of pathogen (specific pathogen-free mice) in the lumen the movement of IELs was reduced compared to that of the infected animals. It is difficult to compare our immunohistochemical study to the live cell imaging reported in the work of Hoytema van Konijnenburg and colleagues. While more CD3^+ cells were associated with fewer F4^+ in the control diet-fed pigs and a similar association was not observed in the yeast-fed pigs, a detailed investigation of the dynamics of IEL CD3+ cells in the small intestine during ETEC infection was not performed. It was also beyond the scope of this work to examine the distribution of T cell subpopulations within in the epithelium. It would be interesting to elaborate our preliminary findings to perform a more detailed characterisation of the IEL CD3^+ cells using this infection model.

The gut microbial ecology findings suggest that the pigs may have developed valuable traits after the exposure to the yeast diet and the bacterial challenge. The divergence of gastrointestinal microbiomes over the course of the ETEC infection was quite distinct for pigs fed either the control or yeast diet. On the second day after the ETEC challenge, the small intestine microbiomes of the yeast fed piglets were more diverse with a co-occurrence between *L. johnsonii* and *L. reuteri*, and *L. reuteri* and *L. mucosae*. In addition, *L. reuteri*, *L. mucosae*, and *L. salivarius* were differentially abundant in the yeast fed pig ileum microbiomes on the second day after the ETEC challenge. No major differences in the large intestine microbiomes were identified on the same day. An exception was higher relative abundance of *Str. luteciae* which was present across the ileum, caecum, and colon microbiomes of piglets fed yeast compared with that of the control-fed piglets. The data obtained by Yang and co-workers suggested that *Str. luteciae* was one of the bacterial phylotypes that was more predominant in the healthy piglet faecal microbiomes compared with those of the piglets with diarrhoea [35]. We could not test this trend on our data since the diarrhoea scoring was performed at a group level.

The transition of the gut microbiomes of piglets fed the yeast diet from d2 to d14 PI was characterized by an increase in alpha diversity of the small intestine microbiome compared with those of the control fed piglets. While various phylotypes increased in numbers in the small intestine, the caecum and colon microbiomes of pigs fed the yeast diet were distinct from those of the control diet on d7 PI. A marked drop in a number of bacterial phylotypes, including various dietary fibre degraders (Figure 7, Figure 8), on d7 PI in the yeast-fed piglet large intestine microbiomes coincided with the loss of co-occurrence of *L. reuteri* and *L. mucosae* in the ileum microbial networks. Interestingly, a decrease in *E. coli* coincided with a decrease on d 7 PI in *L. mucosae* in the ileum of piglets fed yeast compared with that of d2 PI. This may suggest that the clearance of the pathogen by the immune system also targeted *L. mucosae*. In contrast, the populations of host-adapted *L. reuteri* and *L. johnsonii* [36, 37] were neither changed in size nor was their co-occurrence pattern disturbed.

When the co-occurrence of *L. mucosae* and *L. reuteri* was re-instated on d14 PI, the caecum, but not the colon, microbiomes of the piglets fed the yeast diet became more balanced in terms of the differentially abundant phylotypes (Figure 7, Figure 8, Figure 9). The presence of the lactobacilli co-occurrence cluster was another distinct feature of the ileum microbial communities of the yeast-fed piglets. This sub-community was more pronounced in the yeast-fed piglet microbiomes. This distinction in microbial communities may be attributed to the principal differences in the feed formulation. Intact *C. jadinii* yeast cells were fed to animals that cannot enzymatically break down the yeast cell wall components (chitin, mannan-proteins, and yeast beta-glucans). To our knowledge, the ileal digestibility of the yeast feed proteins in weaner piglets is on a par, or higher than that of the proteins from control diets [26, 31]. This means that yeast cell wall disruption is necessary to make yeast intracellular nutrients available for host degradation/uptake. We previously showed that there were more lactobacilli in the small intestine of the yeast-fed healthy piglets compared with that of the control-fed piglets [21]. In the present study, we have also found higher lactobacilli in the ileum and co-occurrence of *L. reuteri* and *L. johnsonii*, and *L. johnsonii* and *L. mucosae* in the yeast-fed piglet gut microbiomes. This consistency in identifying more lactobacilli in the small intestine of piglets fed yeast identifies these bacteria as suitable candidates that are instrumental in degrading yeast cell walls. Tannock et al. demonstrated that *L. johnsonii* and *L. reuteri* could co-exist *in vitro*, and in the mouse forestomach. Also, the authors showed that the two strains could adapt nutrient utilization mechanisms depending on whether the strains were alone or in a co-culture. These two lactobacilli strains can degrade mono- and oligosaccharides via several alternative pathways [38, 39]. However, to degrade complex carbohydrates, the bacteria may be obliged to act in concert to maximize nutrient utilization. *In-silico* analysis of a published porcine gut metagenome database [40] shows that *L. johnsonii* can produce mannan endo-1,4-beta-mannosidase, while *L. reuteri* seems to lack the gene. This enzyme may be operative in the degradation of the yeast cell wall mannan-protein complex.

Charlet and co-workers demonstrated under laboratory conditions that *L. johnsonii* was able to inhibit the growth of live *Candida glabrata* and *Candida albicans* by exerting a chitinase-like activity [41]. The analysis of porcine metagenomic assemblies [40] revealed that both *L. johnsonii* and *L. reuteri* had a gene encoding a LysM domain which is operative in chitin-binding (reviewed in [42]). While both strains can theoretically bind to the yeast cell walls, only *L. johnsonii* seemed to carry chitinase encoding determinants (GH 18). Based on the existing knowledge and our findings, we argue that yeast cells in the feed undergo lactobacilli microbial degradation in the small intestine. We were able to recover a stable connection between *L. johnsonii* and *L. reuteri* from all ileal microbiomes except on d14 PI in the control group using the SPIEC-EASI algorithm.

The two lactobacilli strains are known to be able to colonize non-secretory epithelia and co-exist in biofilms in the alimentary tract of mammals [37, 39].

Based on co-occurrence patterns, our analysis suggests that a distinct lactobacilli phylotype, *L. mucosae*, is the third member of the lactobacilli cluster. As all three strains adhere to surfaces and form biofilms [37, 43], we speculate that these lactobacilli cooperate in degrading the yeast cell wall. In support of this notion, *L. mucosae* was never connected to *L. johnsonii* in the microbiomes of piglets fed diets that did not contain the yeast cell substrate. To pursue this notion further, the microscopy of gastrointestinal tract digesta with lactobacillus species-specific labelling may be useful. Our speculation on the possible role of lactobacillus species could be relevant to animal welfare. Lactobacilli are generally thought to be beneficial bacteria of gastrointestinal tract. Since the *C. jadinii* yeast-derived diet can both fulfil nutritional needs of the animals and possibly augment lactobacilli group, the novel yeast diet could enhance the immunity of the animals. In this study, we have demonstrated that yeast-fed piglets showed loss of appetite. This is an evolutionary adaptation that helps animals withstand life-threatening bacterial infections [32, 33].

Although it is beyond the scope of this work to study the mechanism of appetite loss, we do not exclude possibility of a complementary effect of yeast immunomodulatory components and intestinal lactobacilli to play a key role. A higher microbial diversity in the small intestine may indicate higher tolerance levels of gut immunity. We also speculate that higher microbial diversity of the ileal microbiomes and caecal microbiomes at d14 PI were linked. It is conceivable that richer microbial communities at d14 PI in the ileum are a function of evolved immunologic resilience boosted by the immunogenic properties of yeast. However, further studies are needed to clarify this suggested interaction.

Previous studies have provided evidence that the novel yeast-based diet can support healthy piglets. Irrespective of whether the immune modulation by the yeast diet occurs due to the direct stimulation of the immune system by the yeast beta-glucans and mannans or the indirect stimulation via small intestine lactobacilli growth promotion, or both, the present study shows that the novel diet can improve the health of diseased piglets in herds with a PWD history. However, the response to such diets on the farm is not always comparable to that under controlled experimental conditions. Furthermore, the immunomodulatory properties of yeast are dependent on the species of yeast and down-stream processing conditions of the yeast [44]. Future work should investigate the effect of yeast strain and down-stream processing on nutritional value and health beneficial effects of yeast, and also assess the performance of novel yeast diets under field conditions.

## 5 Ethics statement

The animal study was conducted in compliance with the Norwegian Animal Welfare Act 10/07/2009 and the EU Directive on the Protection of Animals used for Scientific Purposes (2010/63/EU). Norwegian Food Safety Authority approved the use of animals under FOTS ID 16510 protocol.

## 6 Acknowledgements

We thank the Bacteriology group at the Department of Food Safety and Infection Biology at Norwegian University of Life Sciences for their support during sampling. We are grateful to the Histology group at the Department of Basic Sciences and Aquatic Medicine at Norwegian University of Life Sciences for their contribution during sampling and histology analysis. We also thank the staff at the Centre for livestock production (SHF) (NMBU, Ås) and Faculty of Veterinary Medicine, the Department of Production Animal Clinical Sciences (PRODMED) for their excellence in managing the animals during this experiment. Lars Johan Kalager is acknowledged for his help and valuable contributions to this study.

## 7 Methods

### 7.1 Isolation and characterisation of the challenge *E. coli*

The bacterial strain was isolated from a diarrhoea sample of a 31 day old weaner (2 days post-weaning) piglet from a farm with a history of post-weaning diarrhoea (PWD) (described below). The isolate was cultured on blood agar followed by a morphological examination. The bacterial strain was identified as a haemolytic *Escherichia coli* positive for F4 fimbrial antigen as per result of F4(K88) F monovalent rabbit antiserum assay (Statens serum institut, Copenhagen, Denmark). Neo-Sensitabs disc-diffusion antimicrobial susceptibility testing assay(A/S Rosco Diagnostica, Taastrup, Denmark) categorized the strain as being resistant to penicillin, fusidic acid, and streptomycin.

The isolate was propagated on blood agar plate at 37 C for 24 hours. DNA was extracted using a phenol-chloroform method (https://www.pacb.com/wp-content/uploads/2015/09/ SharedProtocol-Extracting-DNA-usinig-Phenol-Chloroform.pdf). The short-read sequencing data were obtained from the Norwegian Veterinary Institute Sequencing unit (SEQ-TECH, VI) (Nextera Flex library prep protocol, Illumina MiSeq 300 bp pair-end sequencing). The long-read data were obtained from Nanopore MinION platform (SQK-RAD004 library prep protocol). Short and long sequencing reads were quality filtered using bbduk version 37.48 (BBMap – Bushnell B., 395 https://sourceforge.net/projects/bbmap/) and Filtlong v0.2.0 (https://github.com/rrwick/Filtlong), respectively. A hybrid (short and long reads) whole-genome assembly was obtained with Unicycler v0.4.8 [45]. The sequenced *E. coli* shared 93.21% genome with *E. coli* UMNK88 NC 017641 (99.81 average nucleotide identity) as per the analysis of the assembled genome using MiGA web-server [46]. Virulence genes of the sequenced *E. coli* were identified using VirulenceFinder 2.0 web-server [47] (https://cge.cbs.dtu.dk/services/VirulenceFinder/). Briefly, the isolate carried genes encoding following virulence determinants: K88/F4, EAST1, heat-labile enterotoxin, and heat-stabile enterotoxin II. The assembled genome was deposited in ENA (ERS5259532).

### 7.2 Experimental design

In total, 68 pure Landrace piglets were used in the study. The animals originated from two farms: a) one with a history of recurrent post-weaning diarrhoea (PWD-immune herd, n = 32) and b) one free of PWD problems (PWD-naive, n = 36). Multiparous sows were given “Porcilis Porcoli Diluvac Forte vet.” and “Porcilis Ery Parvo vet.” (MSD Animal Health, both) before farrowing as a routine vaccination procedure. At day 2 postnatal, piglet oral mucosal swabs were collected followed by DNA extraction using QIAamp DNA Mini Kit (QIAGEN, GmbH, Hilden, Germany). The animals were genotypically classified as being either homozygous (n=48) or heterozygous (n=19) susceptible to F4ac bacterial fimbria adhesion to enterocytes by a *muc*4 gene polymorphism test described previously [8]. Briefly, a DNA fragment of the 5’-CACTCTGCCGTTCTCTTTCC-3’), cleaned (NucleoSpin, Macherey-Nagel), and digested with I restriction enzyme. The susceptible allele was considered if 151 and 216 bp digestion fragments were obtained. No digestion indicated the resistant allele. The piglets were weaned on day 28 ± 2 postnatal (average weight of 8.9±1.5 kg) and transported to the research facility unit where the experiment took place. At weaning, piglets were randomly allocated to either yeast weaner diet or control weaner diet blocking by weight and litter. The resulting four groups, Yeast/PWD-immune, Yeast/PWD-naive, Control/PWD-immune, and Control/PWD-naive, were housed in 4 environmentally controlled pens with dry spruce wood chip bedding (1 pen per each group). The bedding material was renewed twice a day. Feed and water were accessible ad libitum at all times. The diet ingredients and chemical composition are given in the supplementary data (Table 1). Piglets were weaned at 28 days of age. After priming to the weaner diets for one week, all piglets were orally inoculated with 10^9^ CFU/ml (in 2 ml of Lysogeny broth) of F4-positive enterotoxigenic *E. coli*. Faecal swab samples were taken and cultured on blood agar plates to control for the shedding of the challenge strain before and after the inoculation. The animals were sacrificed on day 2, 7, and 14 post-infection (PI) followed by sampling.

### 7.3 Sample collection

Faecal score measurements were taken twice a day throughout the experiment. The faecal scoring system was adopted from [48] which ranged from 1 (firm and shaped) to 4 (watery). The faecal scores were calculated as a mean score per pen per day. Feed leftovers were weighted once a day prior to adding a new portion of the feed. Feed intake was calculated as follows:

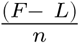

where *F* is the total weight of feed in the feed dispenser on the day before (g), *L* is the weight of leftovers on the current day, (g), and *n* is the number of pigs per pen. Due to the pen level of both faecal scores and feed intake measurements, no statistical procedure was attempted, and the figures were compared directly. Piglets’ body weight was taken at weaning, one-week post-weaning (PW), and at each sampling day for those animals who were euthanised to calculate average daily gain (ADG). ADG was calculated as follows:

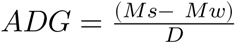

,where *Ms* is weight at sacrifice (kg), *Mw* is weight at weaning (kg), and *D* is the number of days weaning-to-sacrifice (days).

### 7.4 Immunohistochemistry

Formalin-fixed, paraffin-embedded (FFPE) tissues were cut into 4-micron thick sections and mounted on glass slides (SuperFrost Plus, Thermo Scientific™, Braunschweig, Germany) and stored at 4°C until staining. The slides were then incubated at 58°C for 30 min, deparaffinized in xylene and rehydrated in graded alcohols to distilled water. Before the labelling with the primary antibodies, heat-induced antigen retrieval was performed. For immunolabelling with CD3 antibody, the slides were heated in a microwave in Tris-EDTA pH 9.1 buffer with the following steps, first heated to and held at 92°C for 5 min, thereafter the slides were kept in the heated buffer for 5 min. This cycle was repeated with change in the last step where the slides were kept in the heated buffer for 15 min. For immunolabelling with F4 antibody, the slides were heated in an autoclave at 121°C for 10 min in 0.01M, pH6 citrate buffer. Endogenous peroxidase activity was inhibited with 3% H_2_O_2_in methanol for 10 min. Non-specific binding of primary antibody to tissue or Fc receptors was blocked by incubating the slides for 30 min in normal porcine serum at 1:100 in 5% bovine serum albumin (BSA) for CD3 staining and at 1:50 for 20 min for F4 staining. For labelling of T lymphocytes and Fimbrial adhesin F4, monoclonal anti-porcine CD3 clone PPT3 (catalogue number 4510-01, Southern Biotechnology, Birmingham, USA) at 1:1200 and polyclonal rabbit anti F4 (catalogue number 51172, Statens serum institut, Copenhagen, Denmark) at 1:400 were used. The slides were incubated at RT for 1 h, followed by 30 min incubation with secondary antibody. Sections labelled for F4 were incubated with secondary antibody from kit polymer-HRP anti-rabbit (Dako En Vision+ System-HRP, Dako, Glostrup, Denmark) while sections labelled for CD3 were incubated with anti-mouse biotinylated secondary antibody (catalogue number BA-2000-1.5 Vector Laboratories, California, United States) at 1:50 with 1% BSA and thereafter incubated with Vectastain Elite ABC reagent (Vectastain Elite ABC Kit, Vector Laboratories). Detection of peroxidase activity in the F4 and CD3 slides was detected with AEC + substrate from Dako En Vision+ System-HRP and ImmPACT® AEC Substrate, Peroxidase (HRP) (Vector Laboratories), respectively. For counterstaining, hematoxylin was used and as mounting media Aquatex (Merck, Darmstadt, Germany) was used. For enumeration of F4 and CD3 targes, QuPath, v0.2.3 was used (Bankhead2017). The region of interest (ROI) area was determined for F4 and CD3 and used as a reference for quantification: mucosa and the epithelium of four well-oriented villi, respectively. The detection of positive labelling was performed with the following parameters: Gaussian sigma = 2 um, hematoxylin threshold = 0.4, eosin threshold = 0.3. There were three parameters estimated: 1) “F4 counts”, that is the proportion of the total number of mucosal surface-associated F4^+^ *E. coli* positive staining to the mucosa ROI, 2) “F4 size”, that is the average size of the F4+ *E. coli* positive staining areas, or colonies, per the whole area of the section, and 3) “IEL CD3”, that is the proportion of CD3 positive staining per respective epithelial ROI.

### 7.5 Microbial DNA sample handling

At each of the sampling days, 5±1 pigs per pen (12±1 per diet) were euthanised by captive bolt stunning and pithing to allow the collection of gut contents for microbial ecology studies. Digesta from the ileum, caecum, and colon were snap-frozen in liquid nitrogen and stored at -80 C until DNA extraction. Total genomic DNA was extracted from 350 mg of ileum digesta by using QIAamp PowerFecal Pro DNA Kit according to the manufacturer’s instructions, except the samples were homogenized using a bead-beating step with zirconia/silica beads ( =0.1 mm, Carl Roth, Karlsruhe, Germany) in TissueLyser II (Qiagen, Retsch GmbH, Hannover, Germany) with the following parameters: 1) pre-cooling of the TissueLyser adaptors down to 0 C 2) bead-beating 1.5 min at 30 Hz. Total genomic DNA was extracted from 300 mg of the caecum and colon digesta by using QIAamp Fast DNA Stool Mini Kit according to the manufacturer’s instructions, except the bead-beating step described above and digesting proteins with 30 L of Proteinase K II instead of 15-25 L suggested in the manufacturer’s protocol. The purity of extracted DNA was quality controlled by NanoDrop (Thermo Fisher Scientific, Waltham, MA) followed by quantification by Qubit fluorometric broad range assay (Invitrogen, Eugene, OR, USA). Library preparation was performed at the Norwegian Sequencing Centre (https://www.sequencing.uio.no/, Oslo, Norway) using universal prokaryotic primers 319F (5’-ACTCCTACGGGAGGCAGCAG-3’) and 806R (5’-GGACTACNVGGGTWTCTAAT-3’) that amplify the V3-V4 hypervariable region of the 16S *rRNA* gene. Sequencing was performed on a MiSeq sequencer following the manufacturer’s’s guidelines. The resulting demultiplexed raw sequences were deposited in the ENA (PRJEB41033).

### 7.6 Bioinformatics analyses

Demultiplexed paired-end Illumina reads were pre-filtered with bbduk version 37.48 (https://sourceforge.net/projects/bbmap/) by trimming right-end bases less than 15 Phred quality score, removing trimmed reads shorter than 250 bp or/and average Phred quality score less than 20. The resulting reads were further quality filtered by trimming left-end 20 bp and removing reads with maxEE more than 1 for forward and 2 for reverse reads, denoised, merged, and chimera removed with DADA2 R package ver 1.12.1 [49] (Supplementary Figure 10). The resulting ASV tables that derived from two separate Illumina sequencing runs were merged followed by taxonomy assignment using RDP Naive Bayesian Classifier implementation in DADA2 R package (default settings) with GreenGenes database version 13.8, [50] as a reference database. The phylogenetic tree was reconstructed using phangorn R package ver. 2.5.3 [51] under the Jukes-Cantor (JC) nucleotide model with a gamma distribution (k=4, shape=1) with invariant sites (inv=0.2).

DivNet R package [52] was used to estimate Shannon diversity and to test for differences in Shannon diversity estimates in networked gut microbial communities stratified by the day the samples were collected, the gut segment the samples were taken from, with the diet and litter as covariates. Shannon entropy estimator of Phyloseq R package was used to calculate Shannon diversity point estimates. To estimate the Shannon diversity index and to compare it across the microbiomes of the pigs fed distinct diets, DivNet statistical procedure was used for each time point. The beta diversity analysis was performed via the analysis of multivariate homogeneity of group dispersions followed by the permutation test (9999 permutations) on unweighted Unifrac distances and principal coordinate analysis (PCoA) on unweighted Unifrac distances, and permutational multivariate analysis of variance (PERMANOVA) test in R, 9999 permutations. The samples with the read count less than 40000 were discarded from the alpha and beta diversity analyses.

To calculate the relative abundance of bacterial phylotypes in the microbiomes of pigs across diets and time points, group means were taken from the respective groups. To detect differentially abundant bacterial phylotypes, ‘corncob’ algorithm [53] was run on the microbial feature tables (ASV counts per each sample) by fitting a beta-binomial regression model to microbial data for each time point with the diet and litter as covariates. Benjamini-Hochberg correction (cut-off of 0.05) was used to deal with the false discovery rate due to multiple testing. The test was run at each taxonomic level (phylum, class, order, family, species, and ASVs) discarding the samples with the read count less than 10000. Those ASVs lacking genus/species RDP-derived classifications were attempted to be classified manually by using web-based nucleotide BLAST on the non-redundant nucleotide database, where possible. Ambiguous hits were ignored.

### 7.7 Microbial network analysis, ileum

The ASV counts were agglomerated at the genus level and filtered for a minimum of 3 counts per ASV in at least 20% of the samples and at least 50% of the sample per time point (2, 7, 14 days PI) and diet (yeast diet and control diet) using the R package phyloseq version 1.26.1[54]. For each time point and diet, a network was computed on the ileum microbial data with SpiecEasi R package version 1.0.7 [55]. For each recovered network, the edges and nodes were inspected manually.

### 7.8 Statistical analysis

Except otherwise specified, the Bayesian generalized linear models with weakly informative priors were fitted through either bayestesteR v0.7.5 [56] or rstanarm v2.21.1 [57]. The results of the statistical analysis were given as a level of certainty of a certain even to be true given the model and available evidence.

## 8 Supplementary information

**Figure 10:**
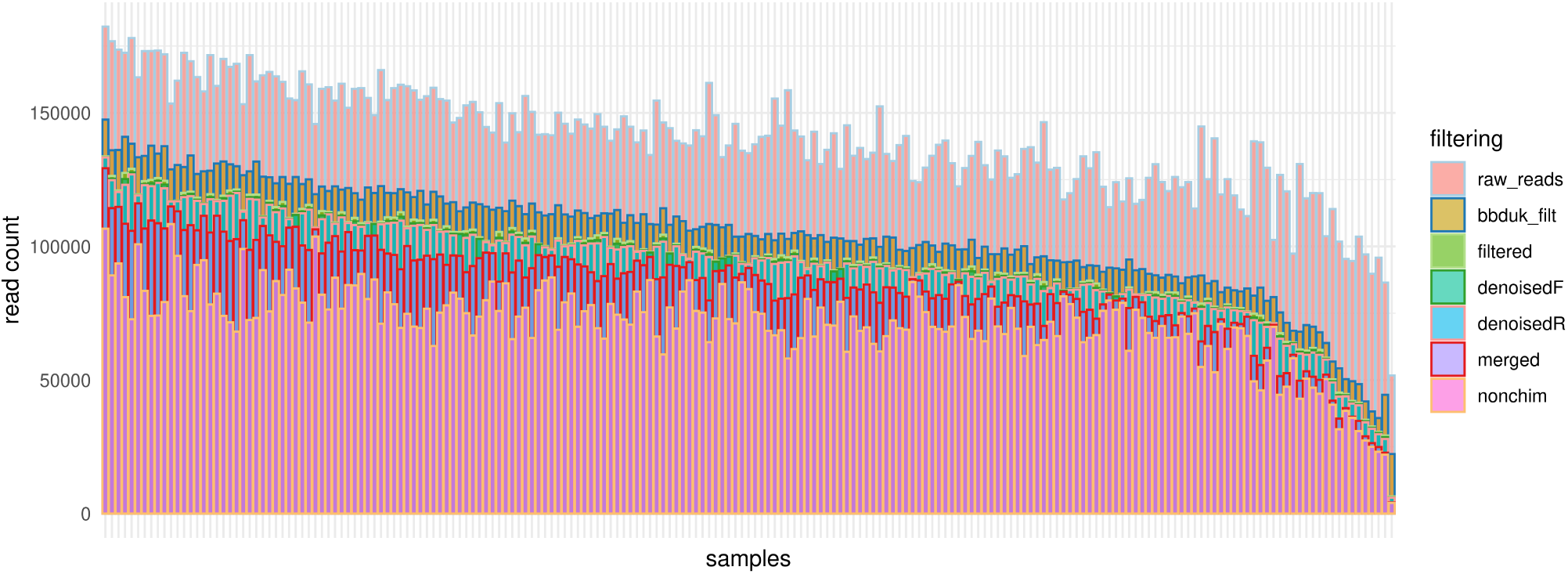
Summary of sequence processing pipeline. The bottom-most bar in the stack (nonchim) shows the number of read that were the basis for making the feature count table (OTU/ASV-table).The bars above nonchim summarise the number of sequencing reads removed at each bioinformatics pipeline step: a) filtered with the bbduk filtering algorithm (*bbduklilt*), b) filtered with the DADA2 algorithm (*filtered*), c) removed after DADA2 denoising step (*denoisedR*/*F*), d) removed due to pair merging failures (*merged*). *rawreads* are raw demultiplexed reads derived from Illumina sequencer.

**Table 1:** Piglet period. Ingredient and chemical composition (g/kg) of diets based on soybean meal (Control) and C. jadinii (Yeast). * Premix : provided the following amounts per kilogram of feed: 120 mg of Zn (ZnO); 460 mg of Fe (FeSO_4_ . H_2_0); 60 mg of Mn (MnO); 26 mg of Cu (CuSO∼4 x 5H_2_O); 0.60 mg of I (Ca(IO_3_)_2_; <1.0 mg of Se (Na_2_SeO_3_); 8000 IU of vitamin A; 1500 IU of cholecalciferol; 45 mg of dl-alpha-tocopheryl acetate; 105 mg of ascorbic acid; 4.64 mg of menadione; 5.63 mg of riboflavin, 3 mg of thiamine; 15 mg of d-pantothenic acid; 20 ug of cyanocobalamine; 45 mg of niacin.

| Ingredients | Control piglet diet | Yeast piglet diet |
| --- | --- | --- |
| Wheat | 627.9 | 593.6 |
| Barley | 100 | 100 |
| Oats | 50 | 50 |
| Yeast meal (*C. jadinii*) (47% CP) | 0 | 146 |
| Soybean meal (SBM) (45% CP) | 80 | 19 |
| Fish meal (68.4% CP) | 20 | 4.8 |
| Potato protein concentrate (72.5% CP) | 33.8 | 9.1 |
| Rapeseed meal (Mestilla) (35%CP) | 20 | 4.9 |
| Rapeseed oil | 19.7 | 23.4 |
| Limestone | 9.2 | 9.4 |
| Monocalcium phosphate | 13.1 | 15.5 |
| Sodium chloride (NaCl) | 7.2 | 5.5 |
| L-Lysine . HCl (98%) | 6.5 | 5.7 |
| L-Threonine | 2.9 | 2.4 |
| L-Methionine | 2.1 | 2.9 |
| L-Valine | 1.4 | 1.2 |
| L-Tryptophan | 0.9 | 0.9 |
| Premix* | 5.3 | 5.5 |
| Calculated contents | - | - |
| Net energy, MJ/kg | 9.94 | 9.94 |
| Crude protein from *C. jadinii*) | 0 | 40 |
| Analyzed content, g/kg | - | - |
| DM | 869 | 885 |
| Gross energy, MJ/kg | 19 | 19 |
| Crude protein | 176 | 172 |
| Crude fat | 39 | 41 |
| Ash | 46 | 45 |
| Neutral detergent fiber (NDF) | 96 | 91 |
| Starch | 442 | 437 |

**Figure 11:**
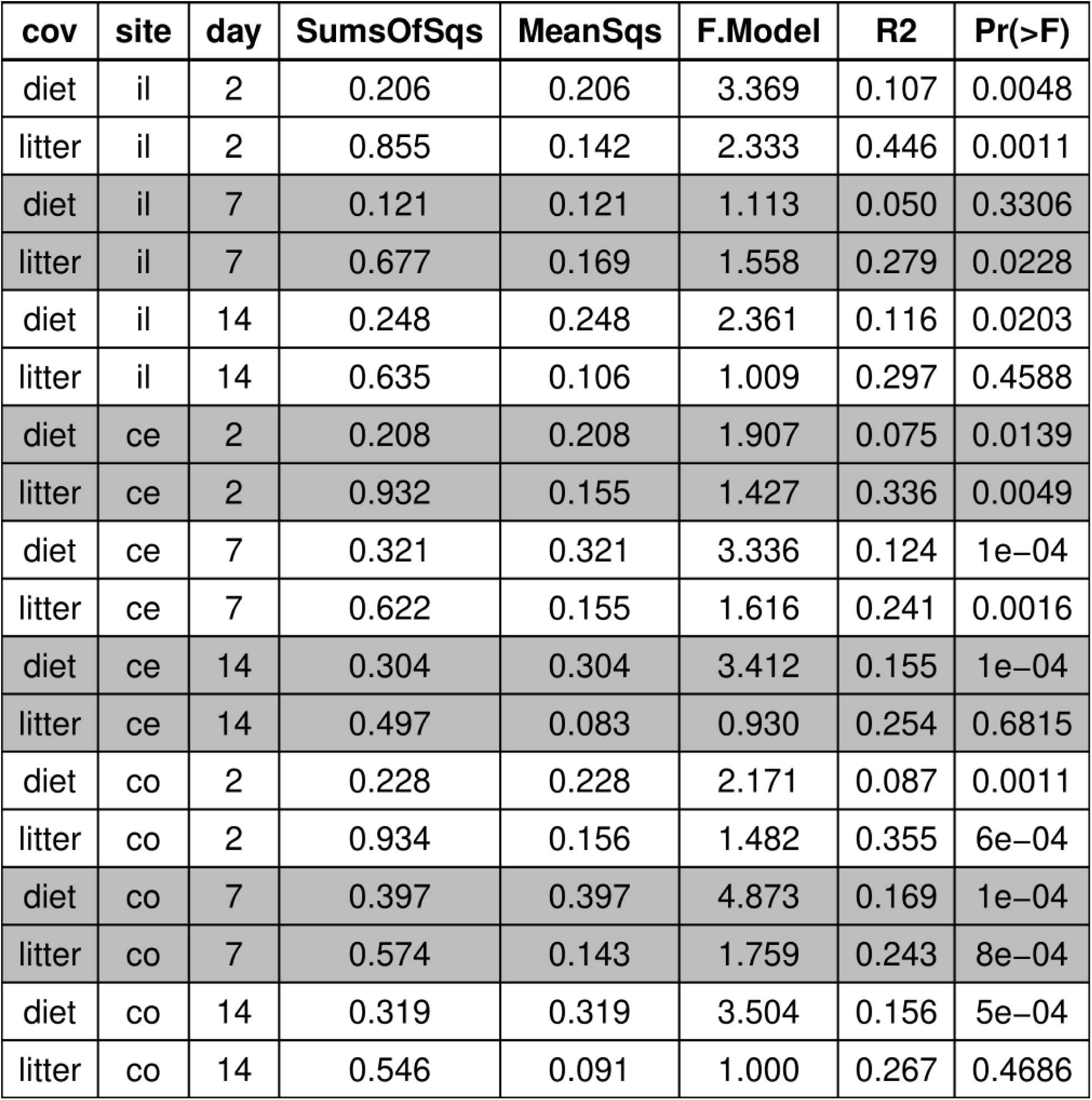
Summary of permutational multivariate analysis of variance (PERMANOVA) test. Each model build on the data across day and gut site is separated by the grey fill.

**Figure 12:**
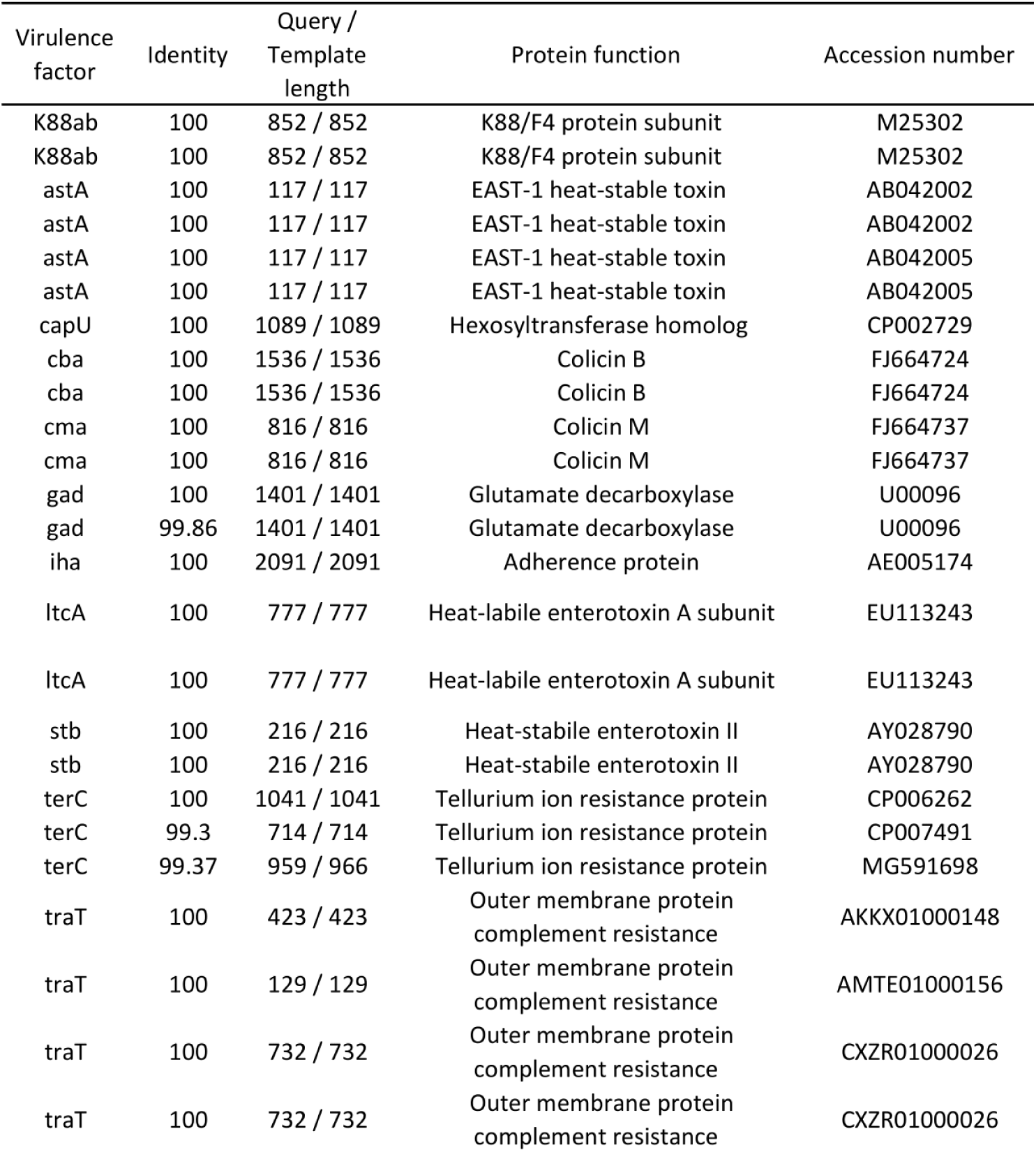
Summary of virulence genes of the *E*. *coli* challenge strain

**Figure 13:**
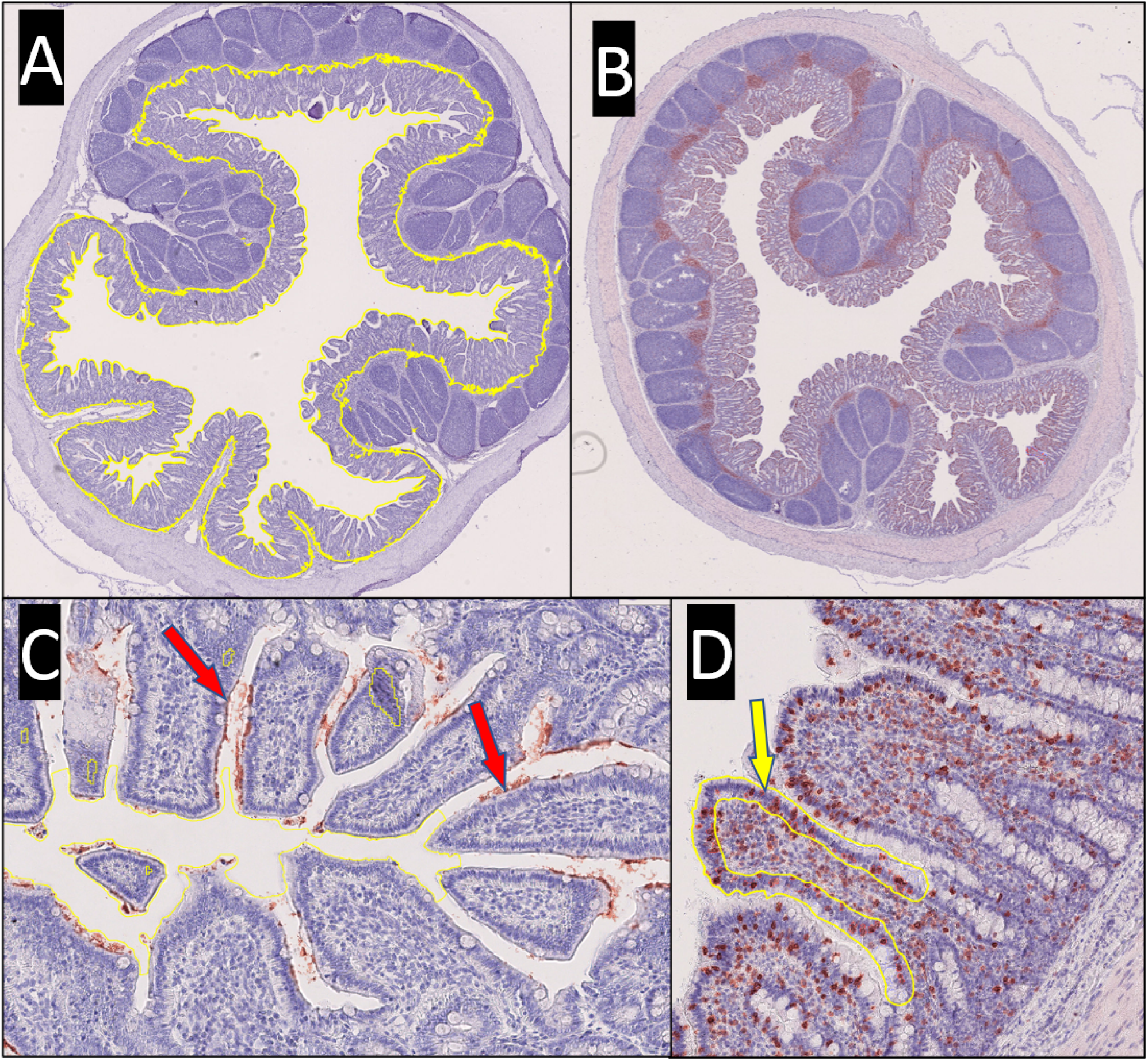
Immunohistochemistry quantification strategy. Panel A: Ileum section with the traced ROI for F4+ quantification. Panel B: Ileum section labelled for CD3+ quantification (hematoxylin). Panel C: Magnification of an ileal section with red arrows pointing at representative F4 colonies that were counted. Panel D: Magnification of an ileal section with the traced ROI for CD3+ quantification.

